## Supplementary material for "Multi-omics analysis in mouse primary cortical neurons reveals complex positive and negative biological interactions between constituent compounds in Centella asiatica": SF1.pdf

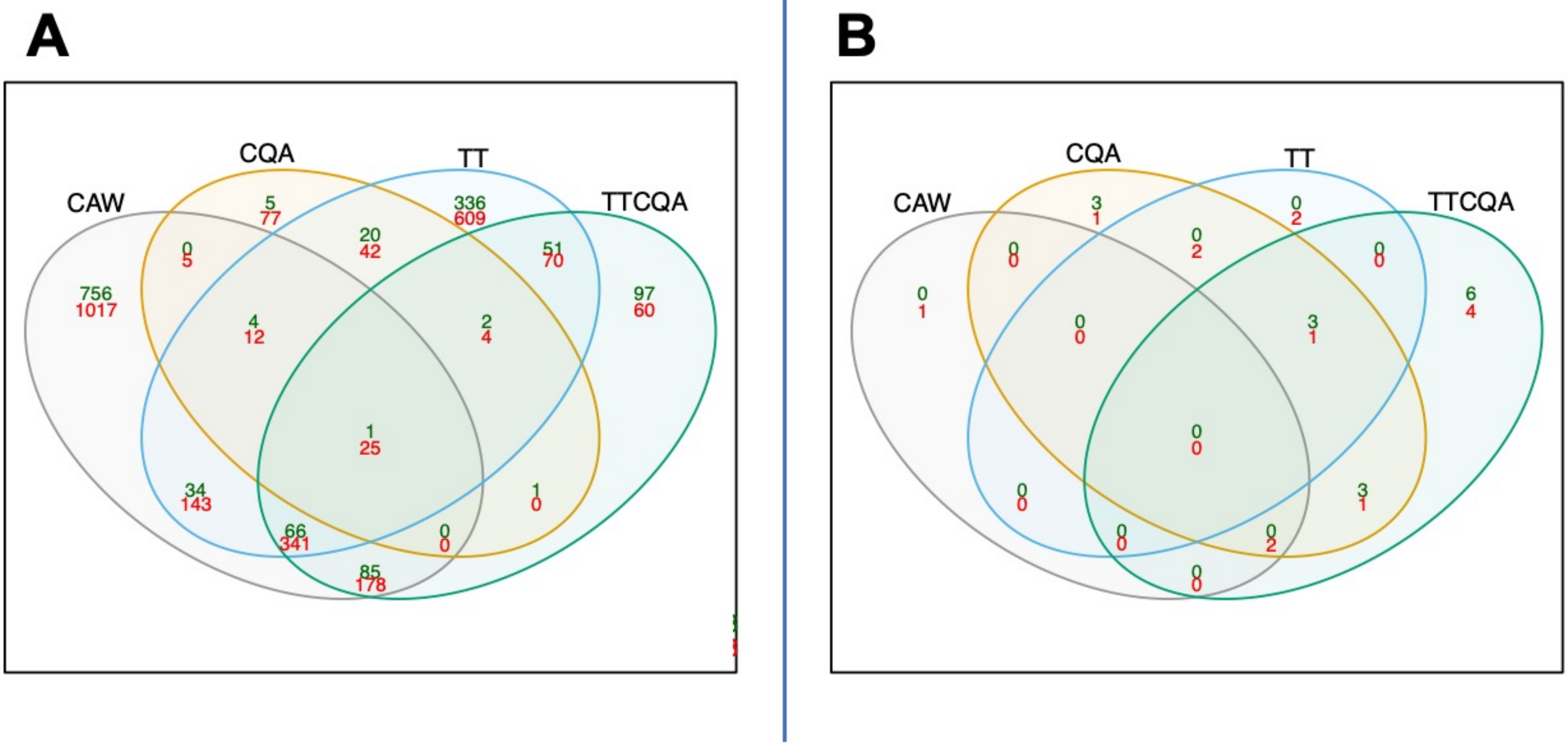


**SF 1. Molecular effects of the four treatments.** Overlap of differential analyses for gene expression **(A)** and metabolite abundance **(B)** for each of the four treatments, relative to control vehicle. The number of upregulated genes/metabolites are shown in green and downregulated genes/metabolites are shown in red.
