## Supplementary material for "Multi-omics analysis in mouse primary cortical neurons reveals complex positive and negative biological interactions between constituent compounds in Centella asiatica": SF2.pdf

**TT Integration Module 1**

(Built using full PPI, using only DE genes, all genes in Module used for pathway enrichment, circled nodes are seeds)

Reactome Pathways:

Metabolism of nucleotides(94/8582,17/19)
Nucleotide salvage(23/8582,10/19)
Purine salvage(14/8582,6/19)
Pyrimidine salvage(10/8582,4/19)
Nucleotide catabolism(35/8582,6/19)
Pyrimidine catabolism(11/8582,3/19)
Nucleotide biosynthesis(14/8582,3/19)
Purine ribonucleoside monophosphate biosynthesis(11/8582,2/19)
Drug ADME(98/8582,6/19)
Azathioprine ADME(26/8582,3/19)
Ribavirin ADME(12/8582,3/19)
Purine catabolism(18/8582,2/19)

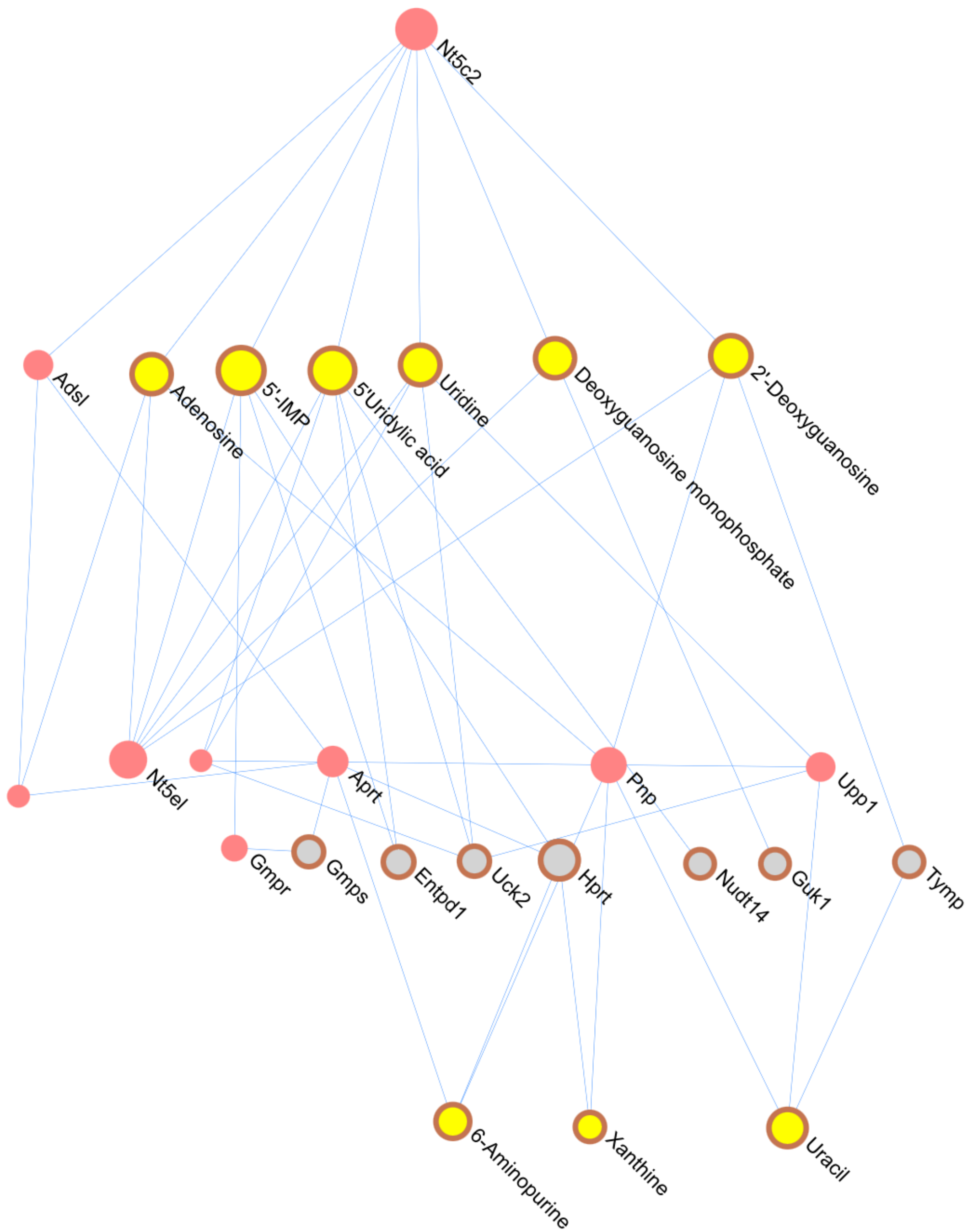

**<u>TT Integration Module 2</u>**

(Built using full PPI, using only DE genes, all genes in Module used for pathway enrichment , circled nodes are seeds)

Reactome Pathways:

Metabolism of amino acids and derivatives(249/8582,14/21)
Aspartate and asparagine metabolism(11/8582,4/21)
Glutamate and glutamine metabolism(13/8582,4/21)
Glyoxylate metabolism and glycine degradation(29/8582,2/21)
Phenylalanine and tyrosine metabolism(12/8582,2/21)
Nucleotide biosynthesis(14/8582,3/21)
Metabolism of nucleotides(94/8582,3/21)
Purine ribonucleoside monophosphate biosynthesis(11/8582,2/21)

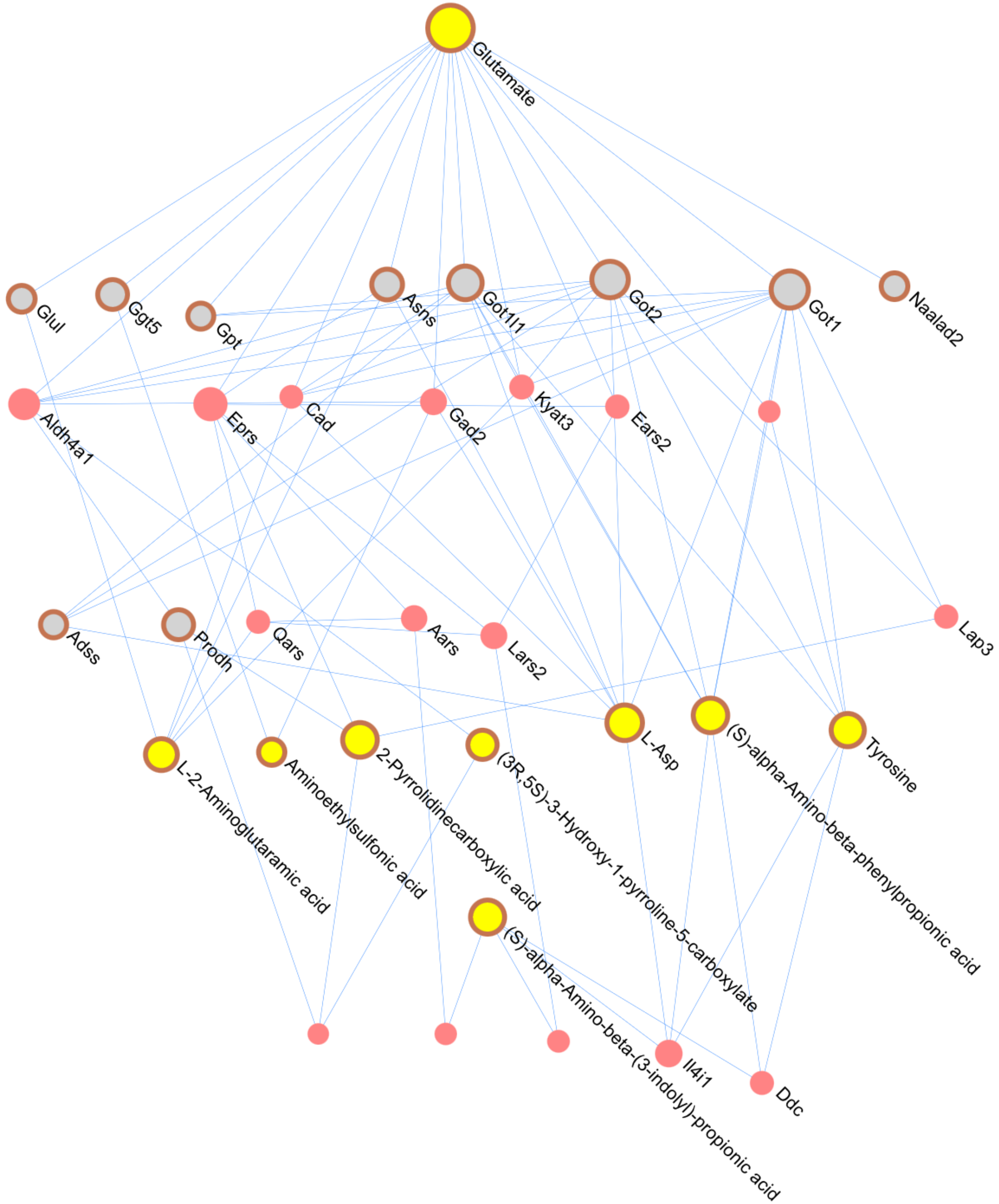

**TT Integration Module 4**

(Built using full PPI, using only DE genes, all genes in Module used for pathway enrichment, circled nodes are seeds)

Reactome Pathways:

Metabolism of amino acids and derivatives(249/8582,9/10)
Metabolism of polyamines(59/8582,3/10)
Protein localization(102/8582,2/10)
Urea cycle(10/8582,4/10)

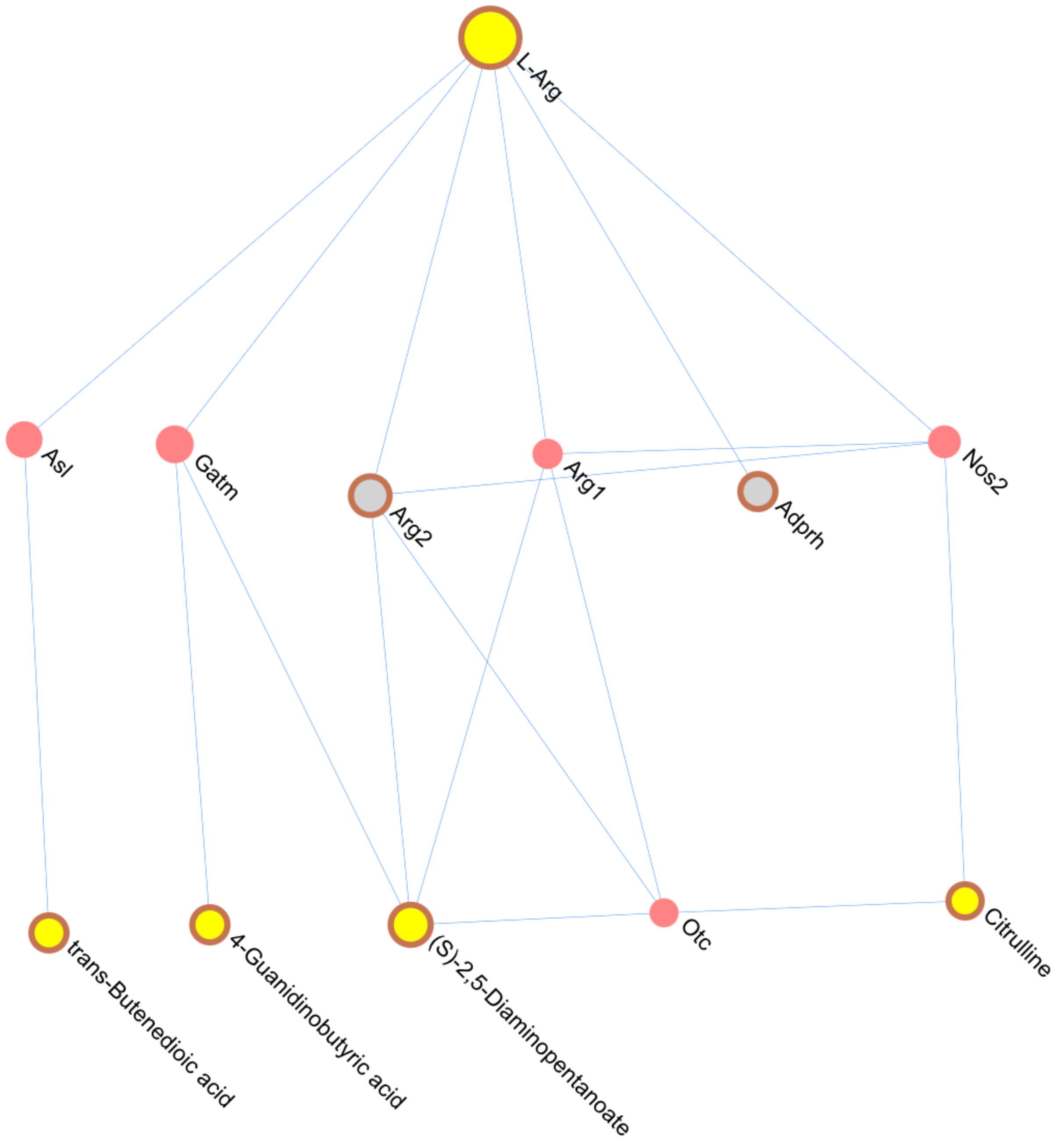

**TT Integration Module 5**
(Built using full PPI, using only DE genes, all genes in Module used for pathway enrichment, circled nodes are seeds)

Reactome Pathways:

Sulfur amino acid metabolism(25/8582,3/12)
Metabolism of amino acids and derivatives(249/8582,4/12)
Methylation(15/8582,3/12)
Phase II - Conjugation of compounds(95/8582,3/12)
Biological oxidations(201/8582,3/12)

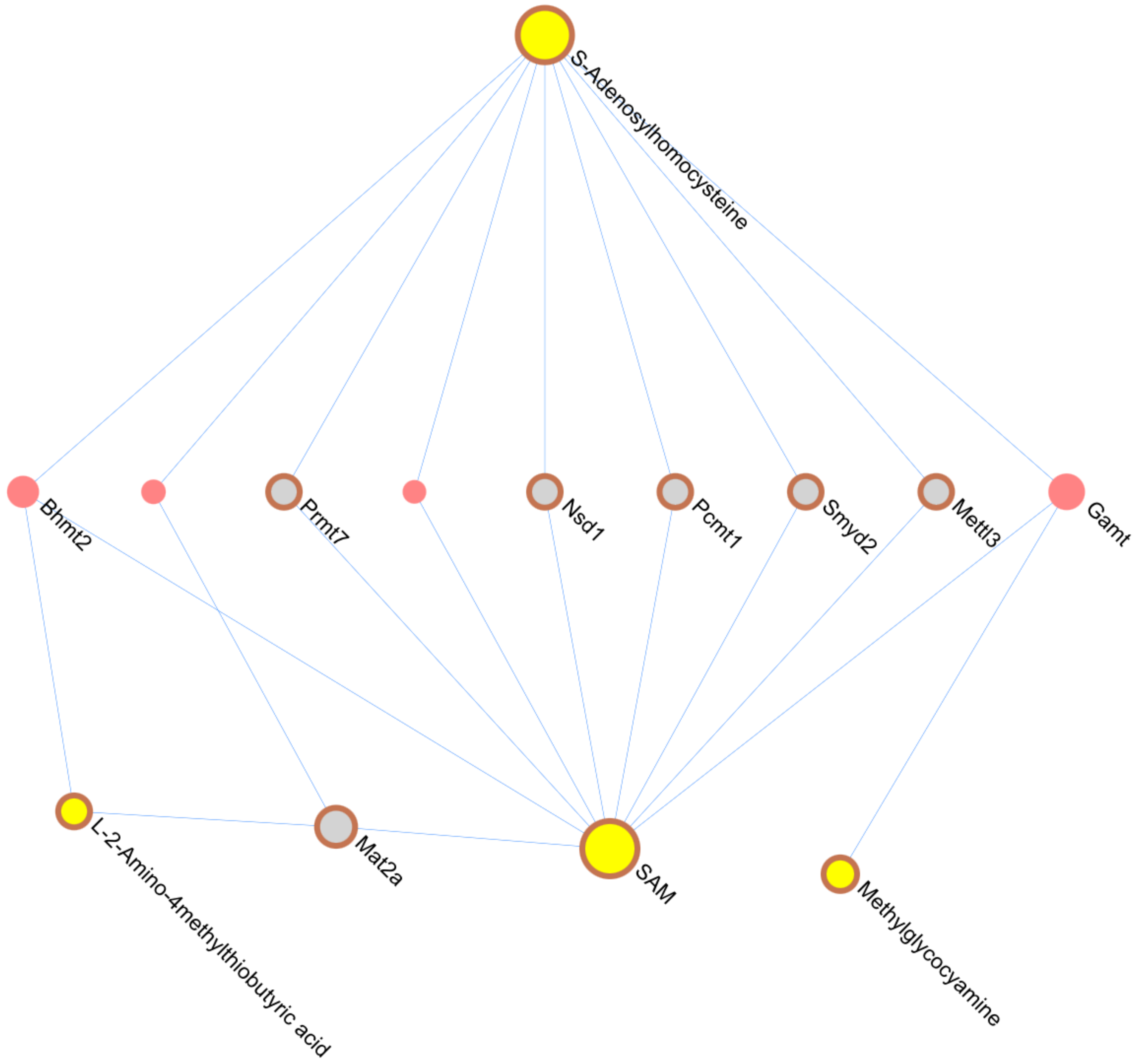

**TT Integration Module 7**
(Built using full PPI, using only DE genes, all genes in Module used for pathway enrichment , circled nodes are seeds)

Reactome Pathways:

Metabolism of amino acids and derivatives(249/8582,3/3)
Branched-chain amino acid catabolism(18/8582,1/3)
Lysine catabolism(12/8582,1/3)

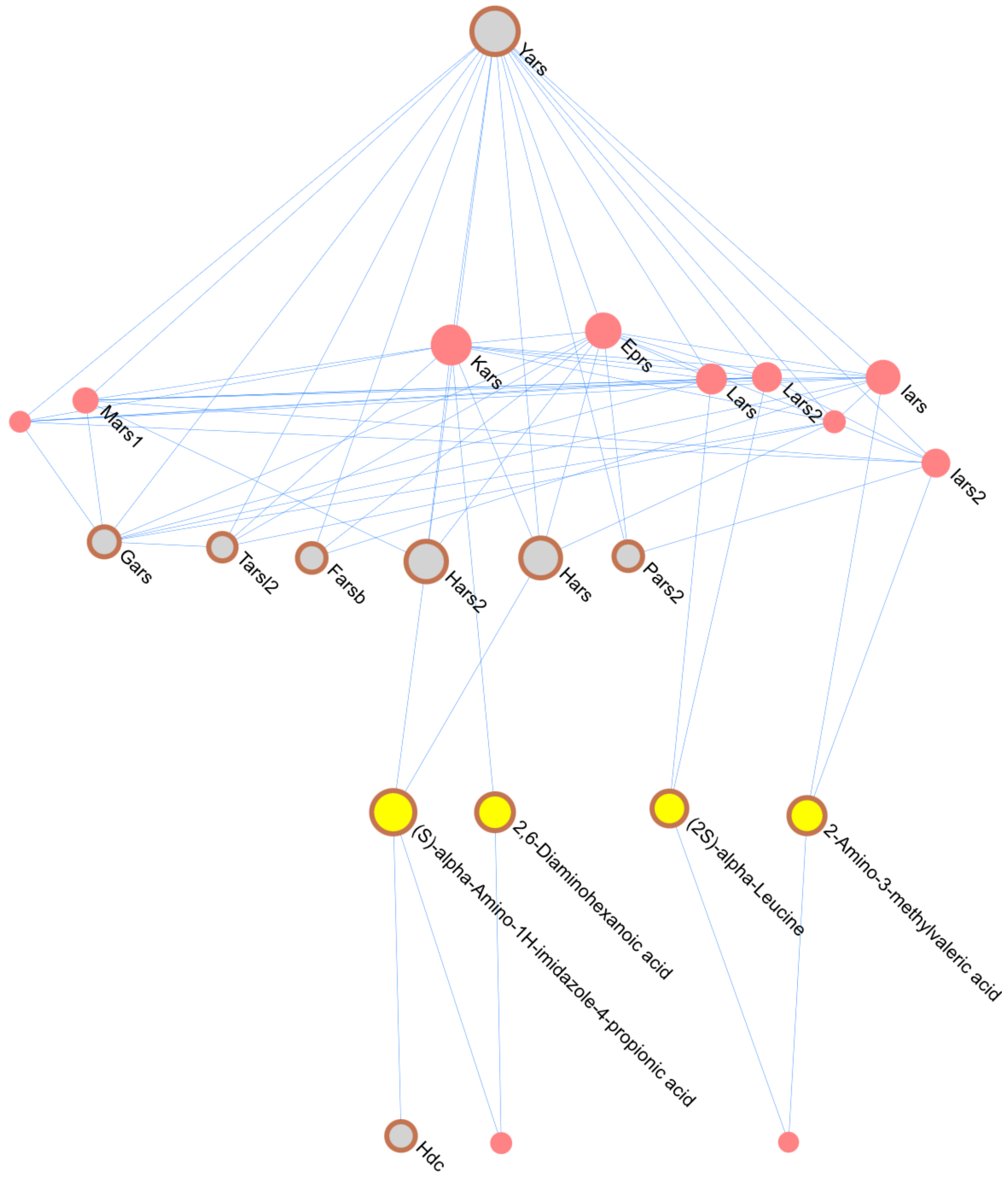

**TT Integration Module 8**

(Built using full PPI, using only DE genes, all genes in Module used for pathway enrichment)

Reactome Pathways:

Fatty acid metabolism(163/8582,6/9)
Arachidonic acid metabolism(53/8582,4/9)
Cytochrome P450 - arranged by substrate type(65/8582,4/9)
Phase I - Functionalization of compounds(101/8582,4/9)
Biological oxidations(201/8582,4/9)
Eicosanoids(11/8582,2/9)
Synthesis of Leukotrienes (LT) and Eoxins (EX)(20/8582,2/9)
Acyl chain remodelling of PI(17/8582,2/9)
Acyl chain remodelling of PS(22/8582,2/9)
Acyl chain remodelling of PC(27/8582,2/9)
Acyl chain remodelling of PE(27/8582,2/9)
Peroxisomal protein import(65/8582,2/9)
Mitochondrial Fatty Acid Beta-Oxidation(36/8582,2/9)

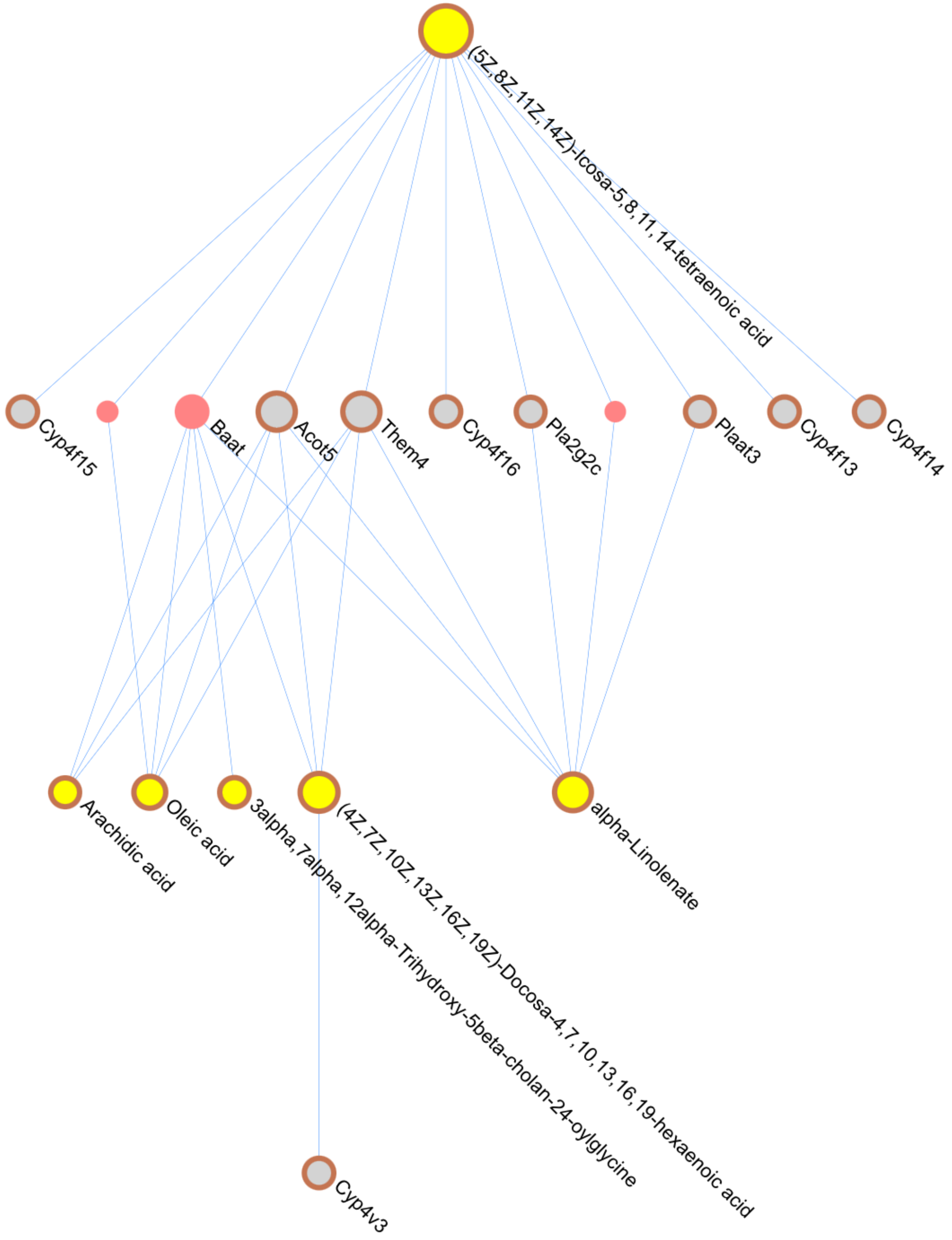
