## Supplementary material for "Multi-omics analysis in mouse primary cortical neurons reveals complex positive and negative biological interactions between constituent compounds in Centella asiatica": SF3.pdf

### Pre and Post Normalized Distributions (1-50)

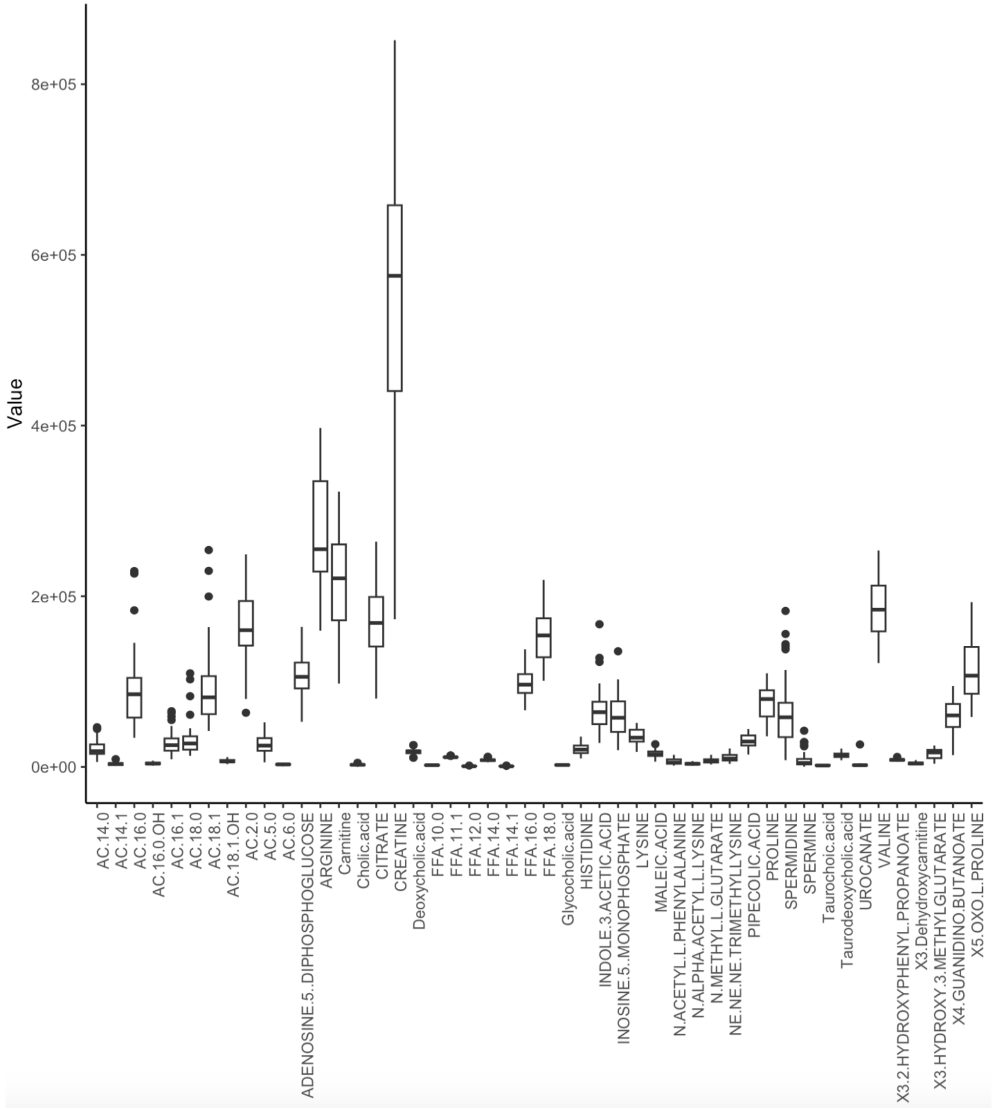


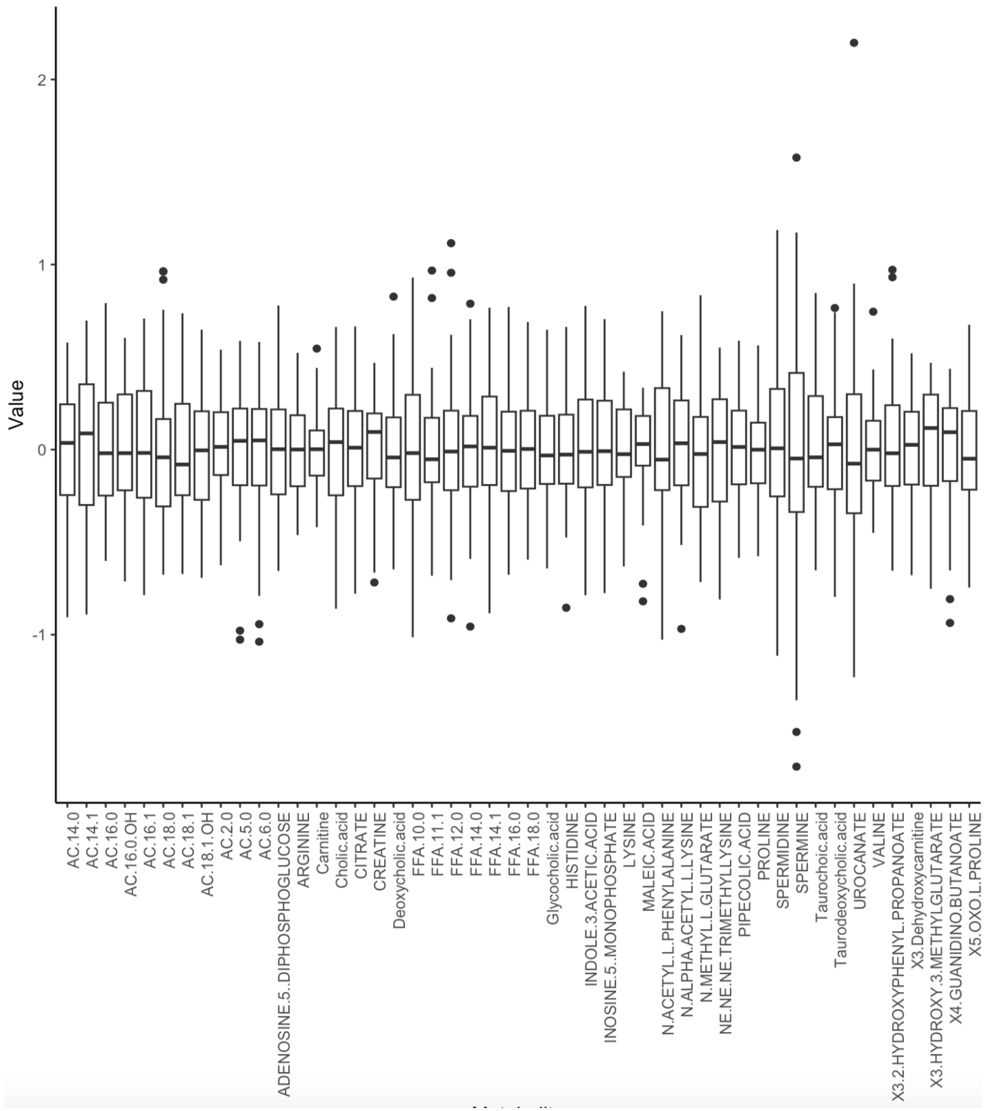

### Pre and Post Normalized Distributions (51-100)

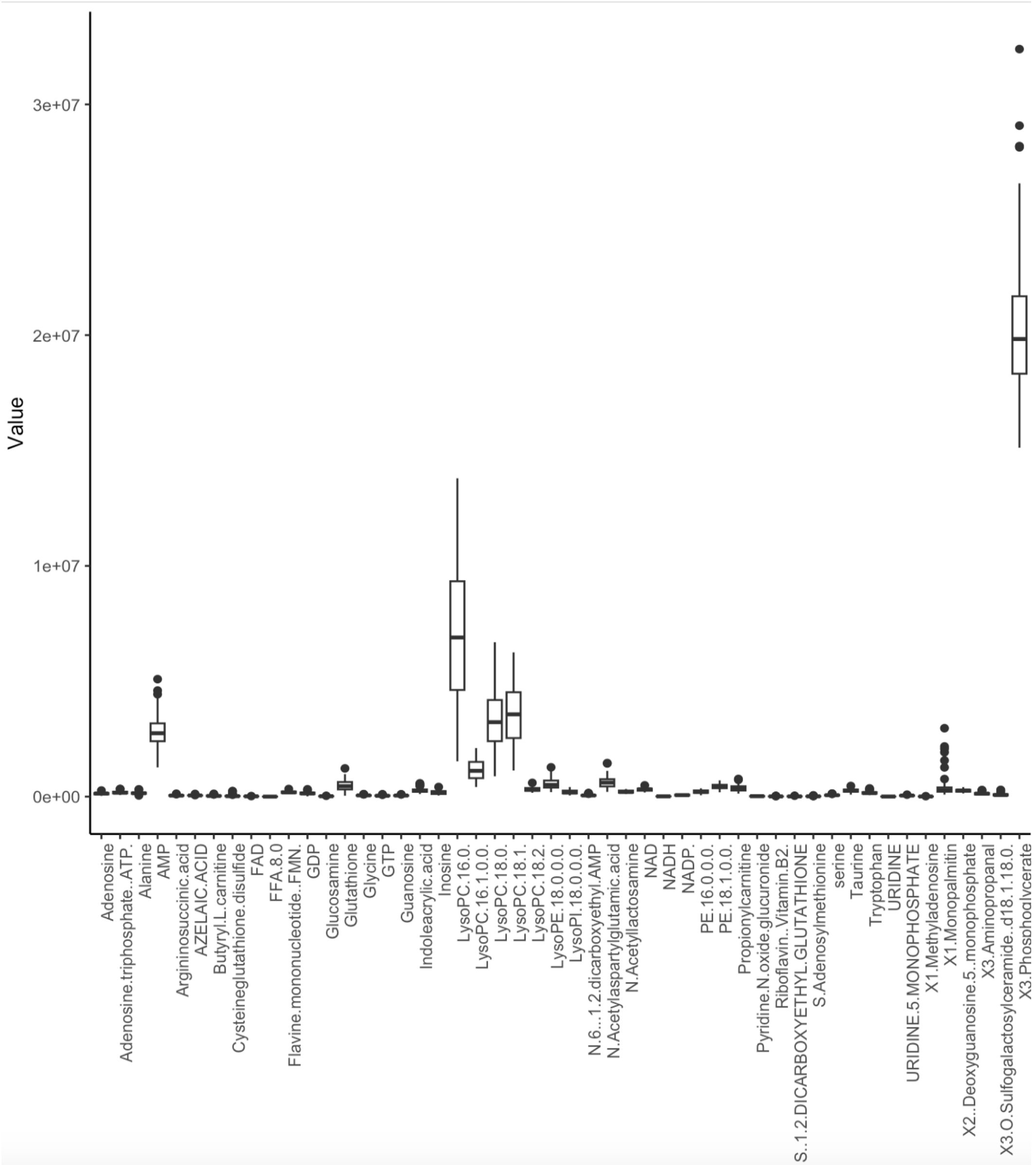


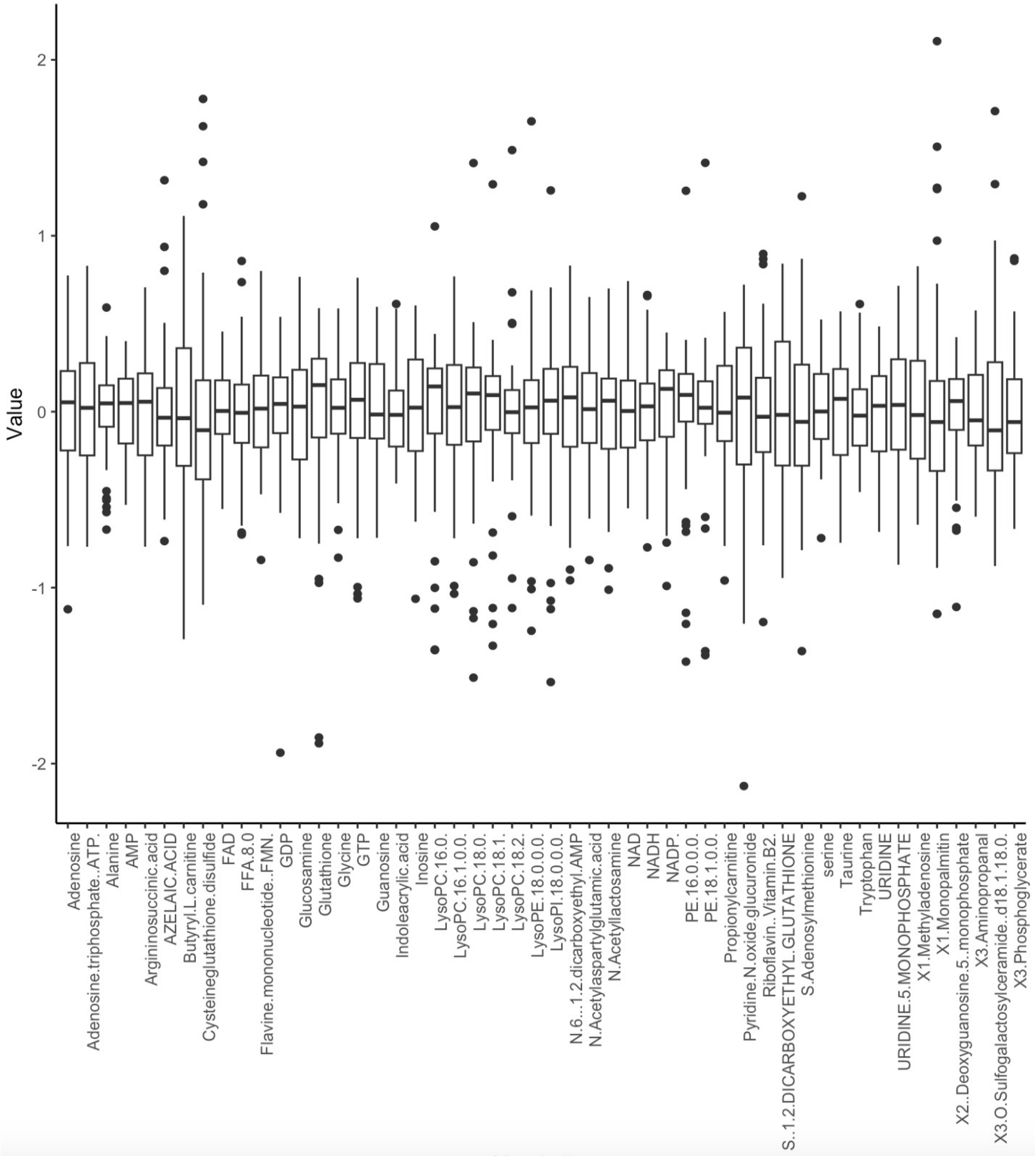

### Pre and Post Normalized Distributions (101-150)

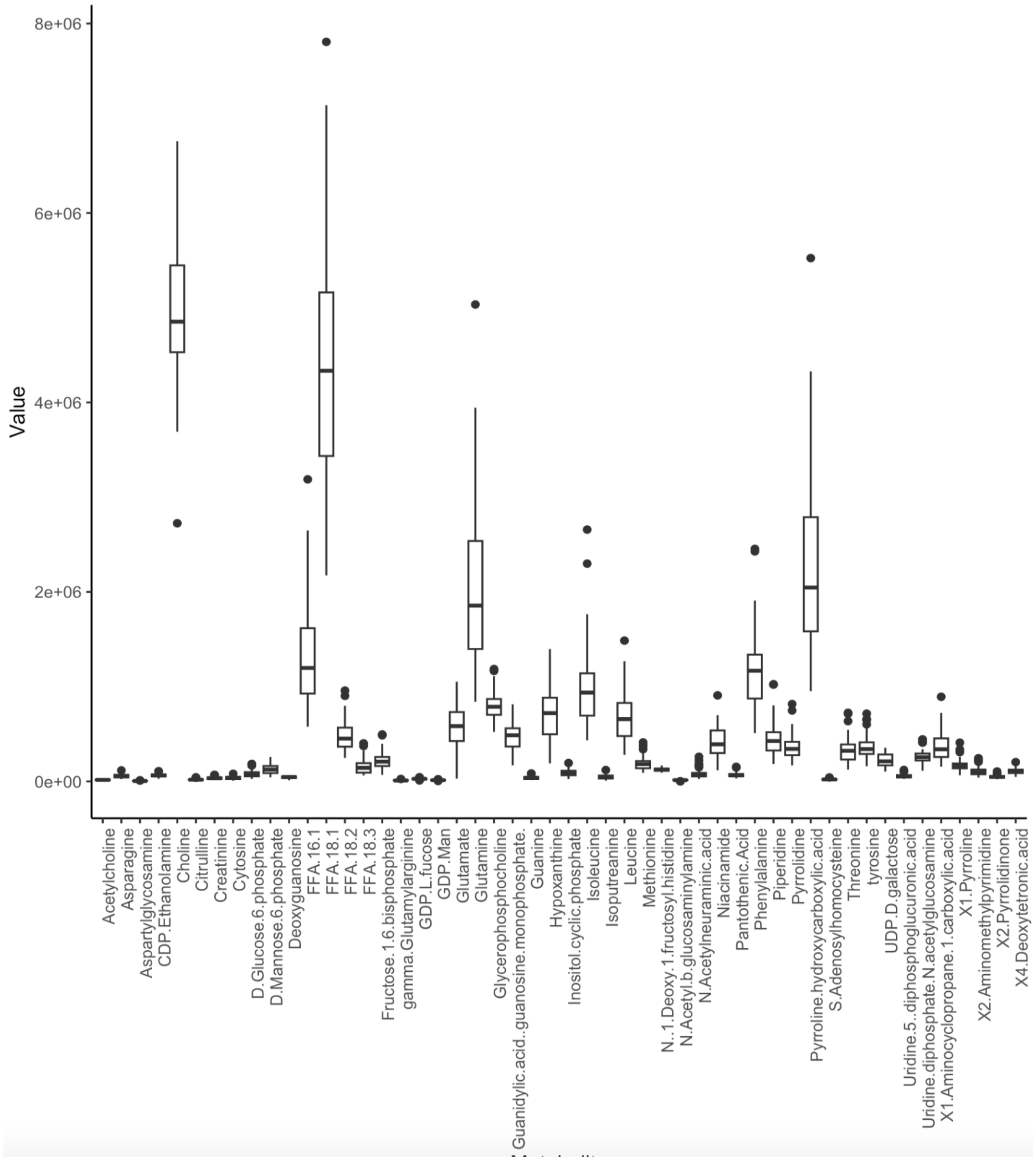


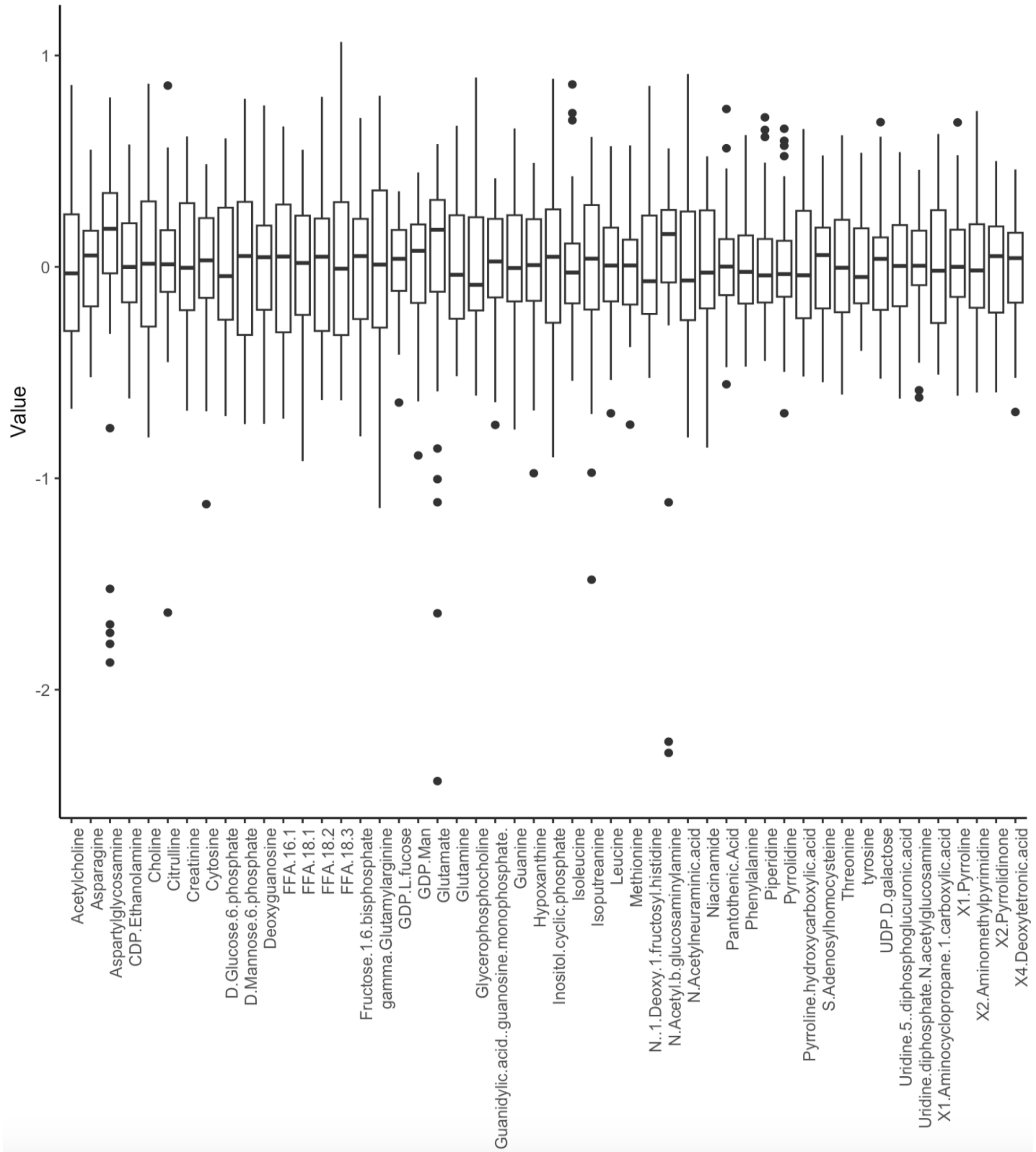

### Pre and Post Normalized Distributions (151-195)

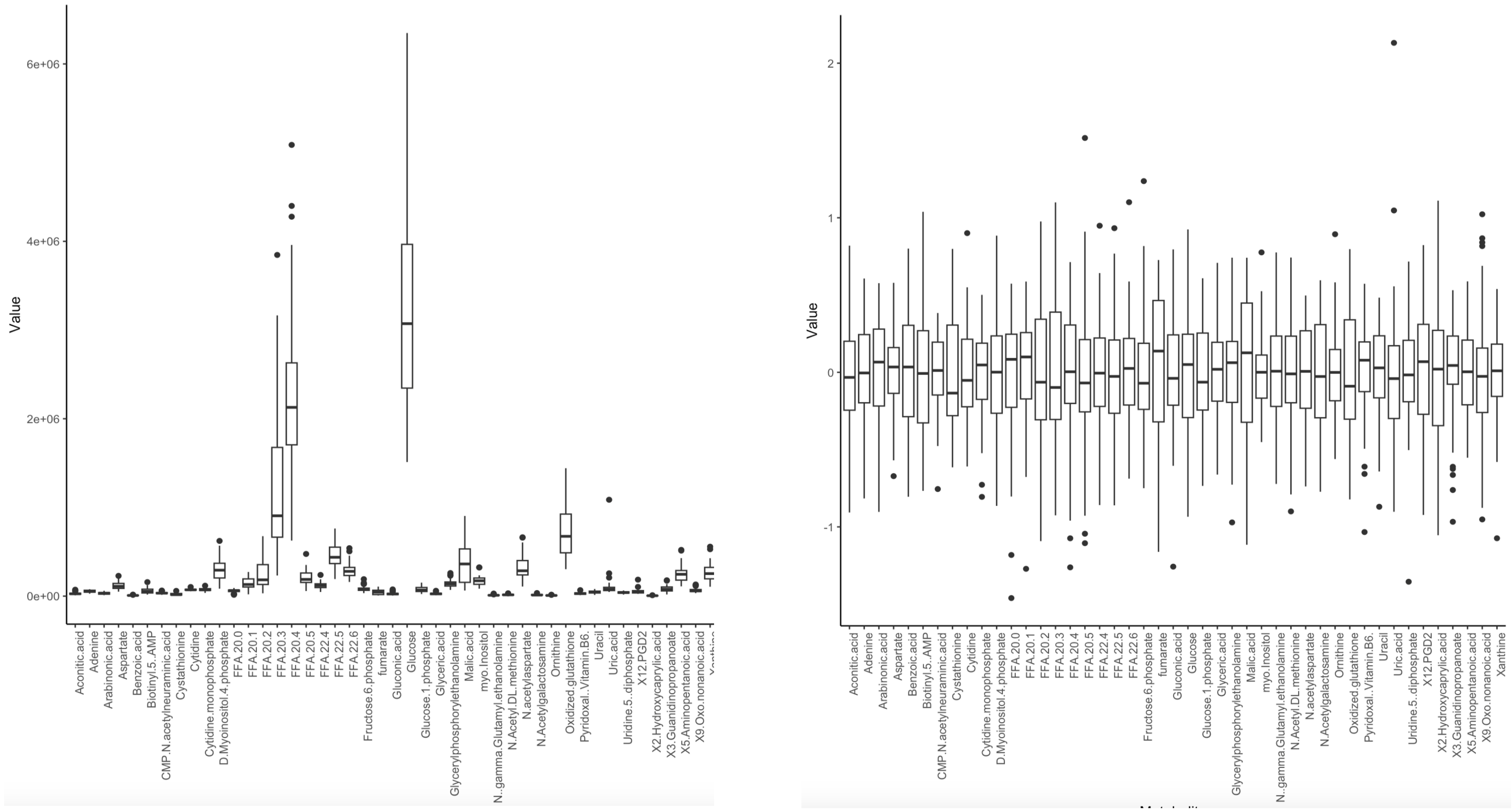
