## Supplementary material for "Multi-omics analysis in mouse primary cortical neurons reveals complex positive and negative biological interactions between constituent compounds in Centella asiatica": SF4.pdf

**CQA integrated module 1:**
(Built using full PPI, using only DE genes, all genes in Module used for pathway enrichment, circled nodes are seeds)

110 pathways

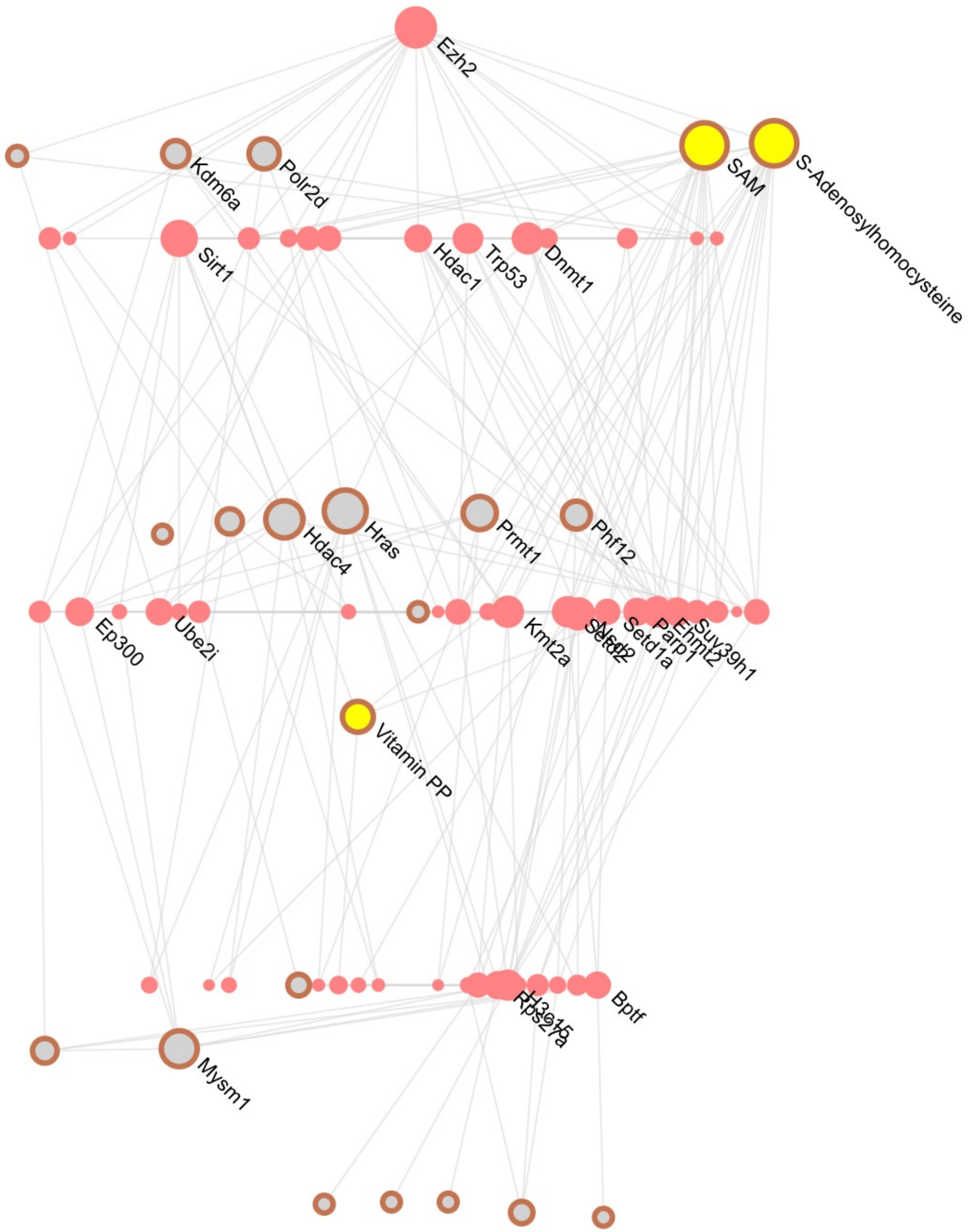

**<u>CQA integrated module 2:</u>**

(Built using full PPI, using only DE genes, all genes in Module used for pathway enrichment, circled nodes are seeds)

Metabolism of nucleotides(94/8582,14/16)
Nucleotide salvage(23/8582,8/16)
Pyrimidine salvage(10/8582,2/16)
Purine salvage(14/8582,6/16)
Ribavirin ADME(12/8582,4/16)
Drug ADME(98/8582,5/16)
Nucleotide catabolism(35/8582,4/16)
Pyrimidine catabolism(11/8582,2/16)
Nucleotide biosynthesis(14/8582,4/16)
Purine ribonucleoside monophosphate biosynthesis(11/8582,3/16)
Purine catabolism(18/8582,2/16)

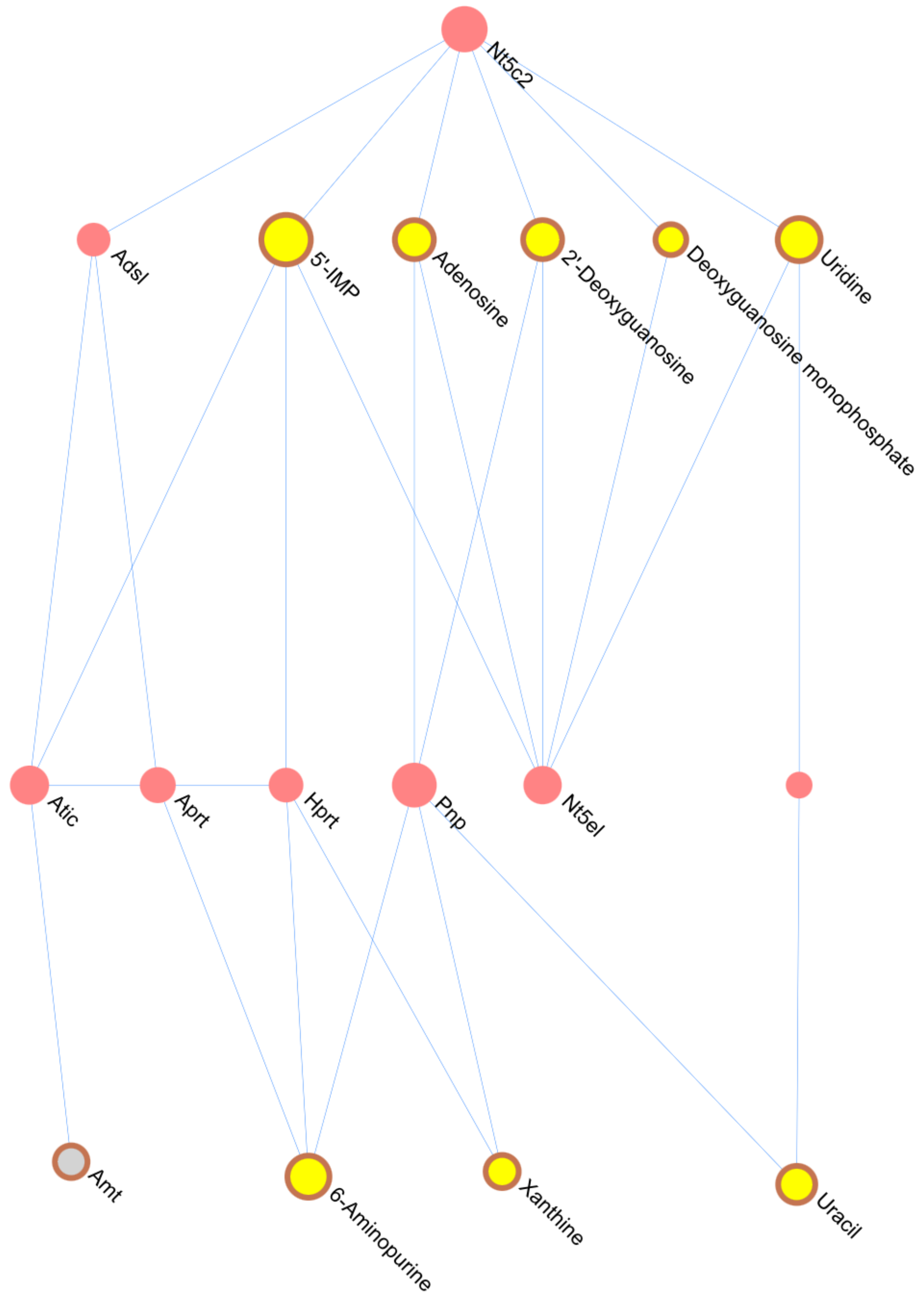

**CQA integrated module 3:**
(Built using full PPI, using only DE genes, all genes in Module used for pathway enrichment, circled nodes ar seeds)

Metabolism of amino acids and derivatives(249/8582,3/3)
Glutamate and glutamine metabolism(13/8582,2/3)

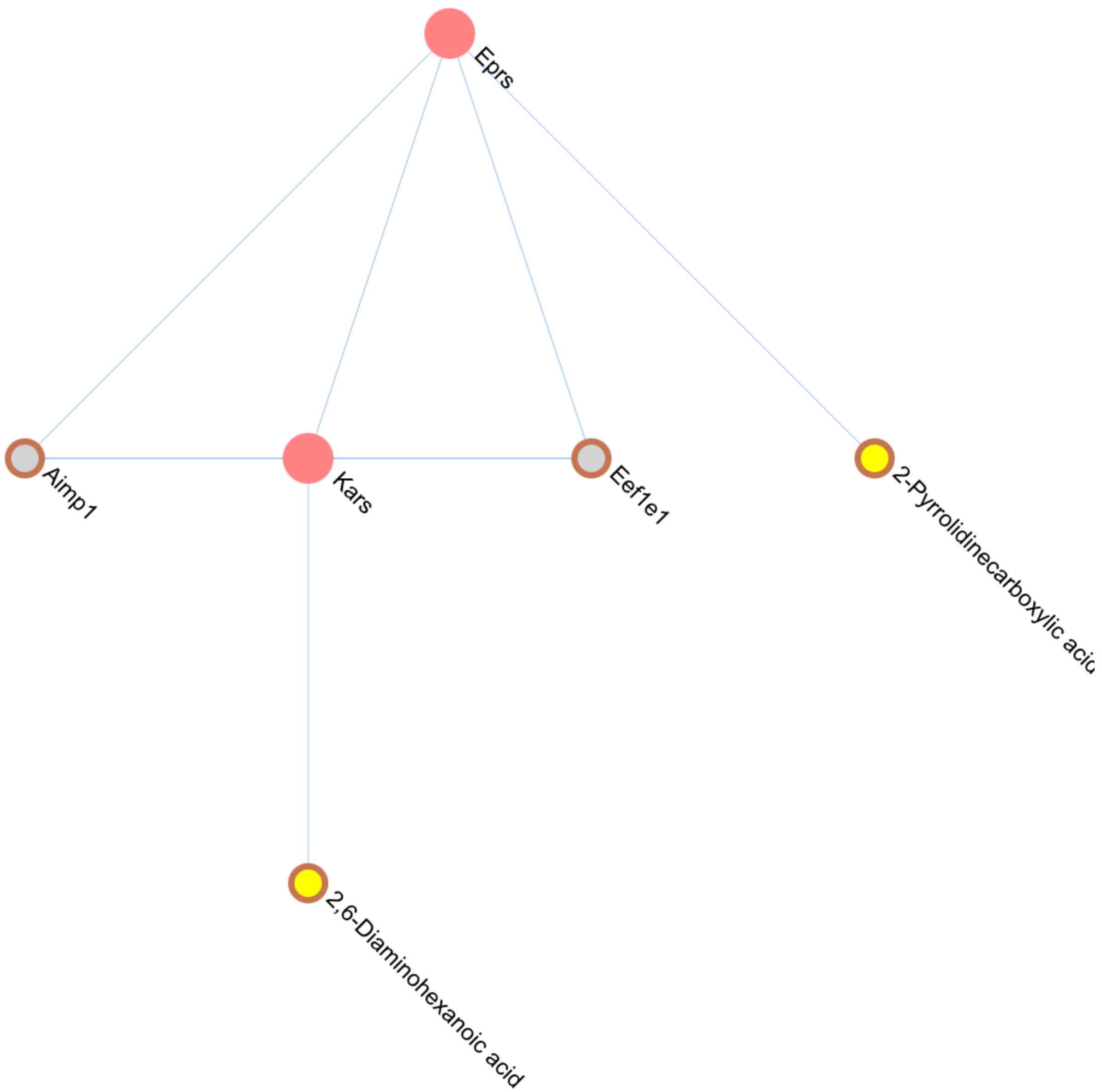

**CQA integrated module 4:**
(Built using full PPI, using only DE genes, all genes in Module used for pathway enrichment, circled nodes are seeds)

Glucose metabolism(80/8582,5/9)
Glycolysis(62/8582,4/9)
Metabolism of carbohydrates(263/8582,5/9)
Transport of Mature mRNA Derived from an Intronless Transcript(40/8582,3/9)
Transport of Mature mRNAs Derived from Intronless Transcripts(41/8582,3/9)
Transport of Mature Transcript to Cytoplasm(79/8582,3/9)
Regulation of Glucokinase by Glucokinase Regulatory Protein(30/8582,2/9)
Processing of Capped Intron-Containing Pre-mRNA(265/8582,3/9)
Processing of Intronless Pre-mRNAs(20/8582,2/9)
Processing of Capped Intronless Pre-mRNA(29/8582,2/9)
mRNA 3'-end processing(55/8582,2/9)
RNA Polymerase II Transcription Termination(64/8582,2/9)

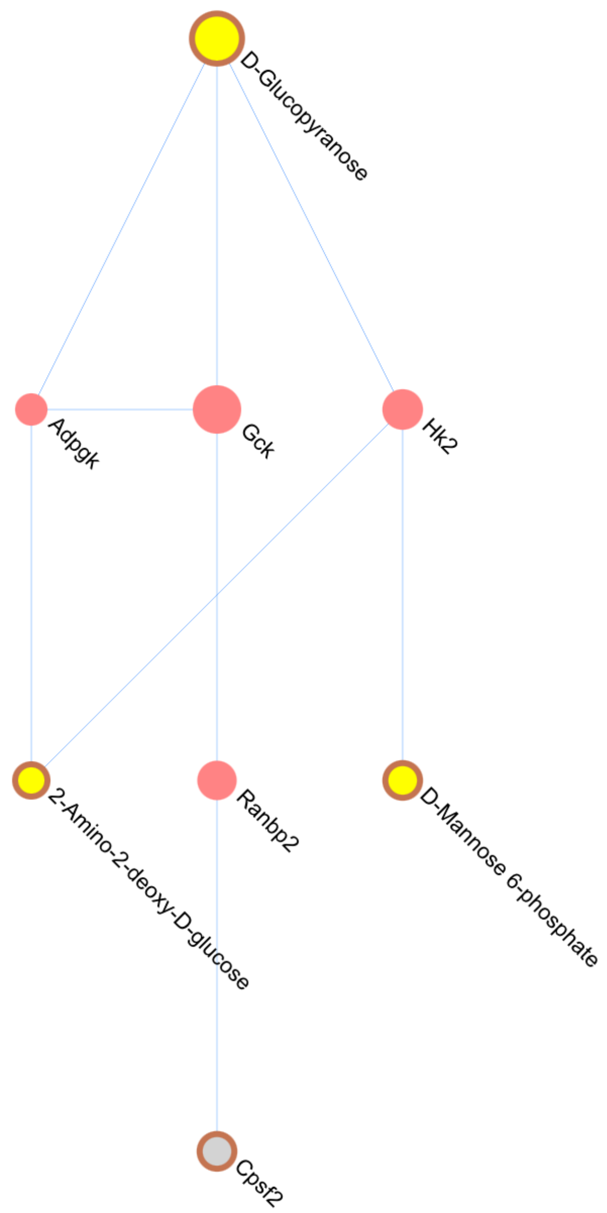

### CQA integrated module 6:

(Built using full PPI, using only DE genes, all genes in Module used for pathway enrichment, circled nodes are seeds)

Metabolism of amino acids and derivatives(249/8582,13/40)
The citric acid (TCA) cycle and respiratory electron transport(163/8582,13/40)
Citric acid cycle (TCA cycle)(21/8582,7/40)
Pyruvate metabolism and Citric Acid (TCA) cycle(47/8582,8/40)
Gluconeogenesis(34/8582,6/40)
Glucose metabolism(80/8582,7/40)
Metabolism of carbohydrates(263/8582,9/40)
Mitochondrial biogenesis(27/8582,6/40)
Transcriptional activation of mitochondrial biogenesis(10/8582,3/40)
Glutamate and glutamine metabolism(13/8582,3/40)
Organelle biogenesis and maintenance(212/8582,6/40)
Pentose phosphate pathway(13/8582,2/40)
Respiratory electron transport, ATP synthesis by chemiosmotic coupling, and heat producti
on by uncoupling proteins.(116/8582,5/40)
Aspartate and asparagine metabolism(11/8582,3/40)
Glyoxylate metabolism and glycine degradation(29/8582,3/40)
Glycolysis(62/8582,4/40)
Methylation(15/8582,2/40)
Urea cycle(10/8582,2/40)
Formation of ATP by chemiosmotic coupling(17/8582,3/40)
Cristae formation(17/8582,3/40)

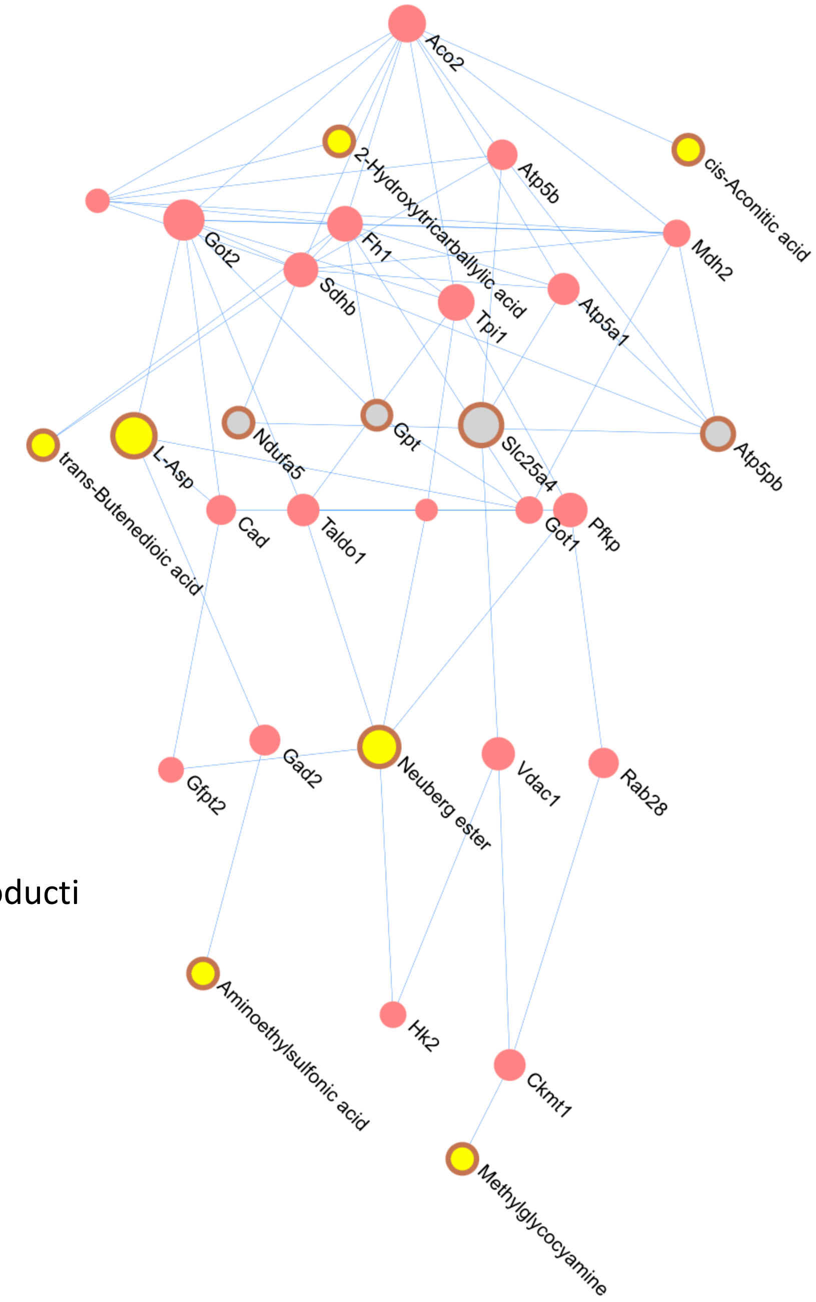
