## Supplementary material for "Multi-omics analysis in mouse primary cortical neurons reveals complex positive and negative biological interactions between constituent compounds in Centella asiatica": SF5.pdf

**TTCQA integrated module 8:**
(Built using full PPI, using only DE genes, all genes in Module used for pathway enrichment, circled nodes are seeds)

91 pathways

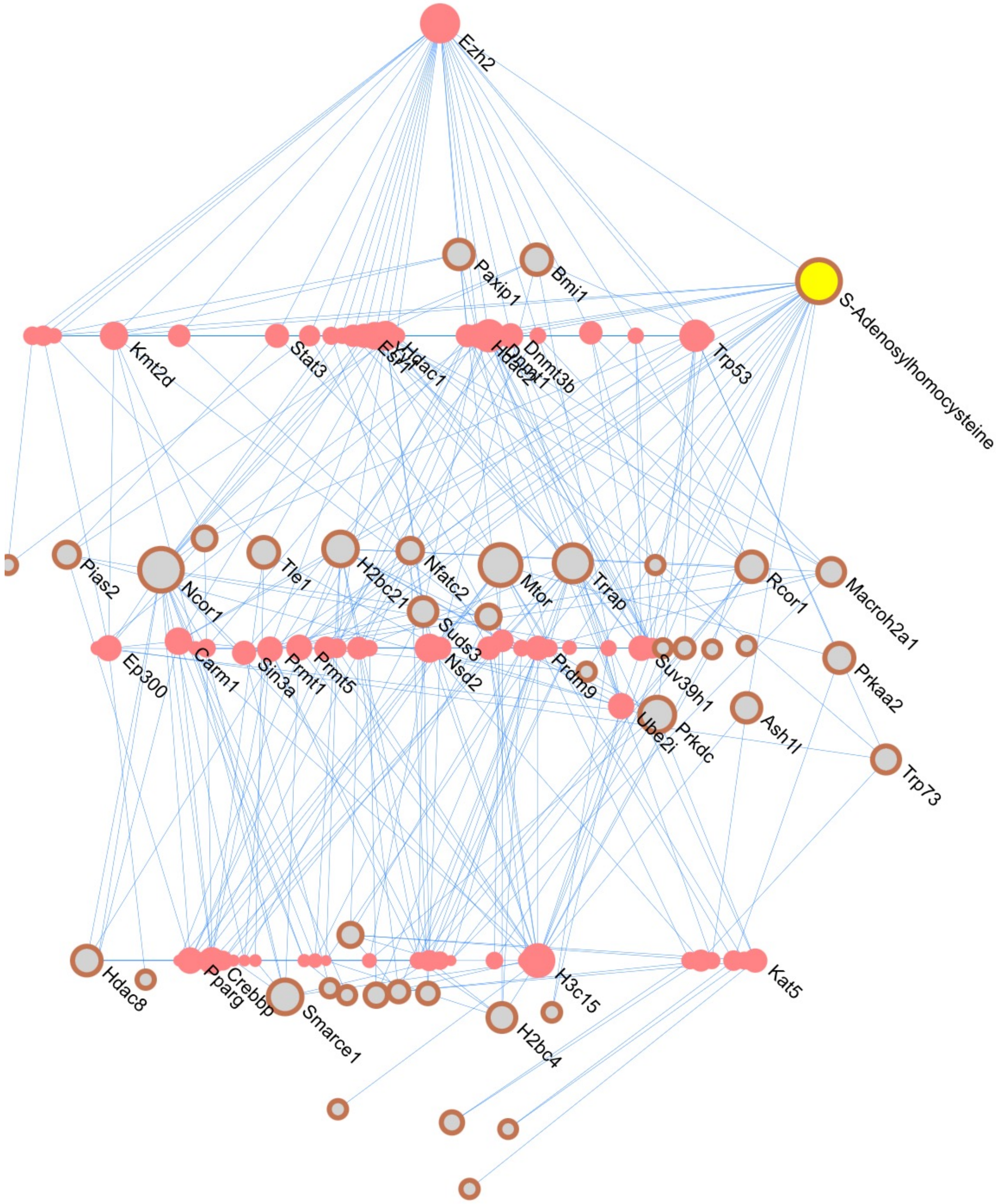

**TTCQA integrated module 16:**

(Built using full PPI, using only DE genes, all genes in Module used for pathway enrichment, circled nodes are seed

Fatty acid metabolism(163/8582,7/11)
Arachidonic acid metabolism(53/8582,5/11)
Cytochrome P450 - arranged by substrate type(65/8582,5/11)
Phase I - Functionalization of compounds(101/8582,5/11)
Biological oxidations(201/8582,5/11)
Endogenous sterols(27/8582,2/11)
Biosynthesis of DHA-derived SPMs(17/8582,3/11)
Biosynthesis of specialized proresolving mediators (SPMs)(18/8582,3/11)
Xenobiotics(26/8582,3/11)
Mitochondrial Fatty Acid Beta-Oxidation(36/8582,2/11)

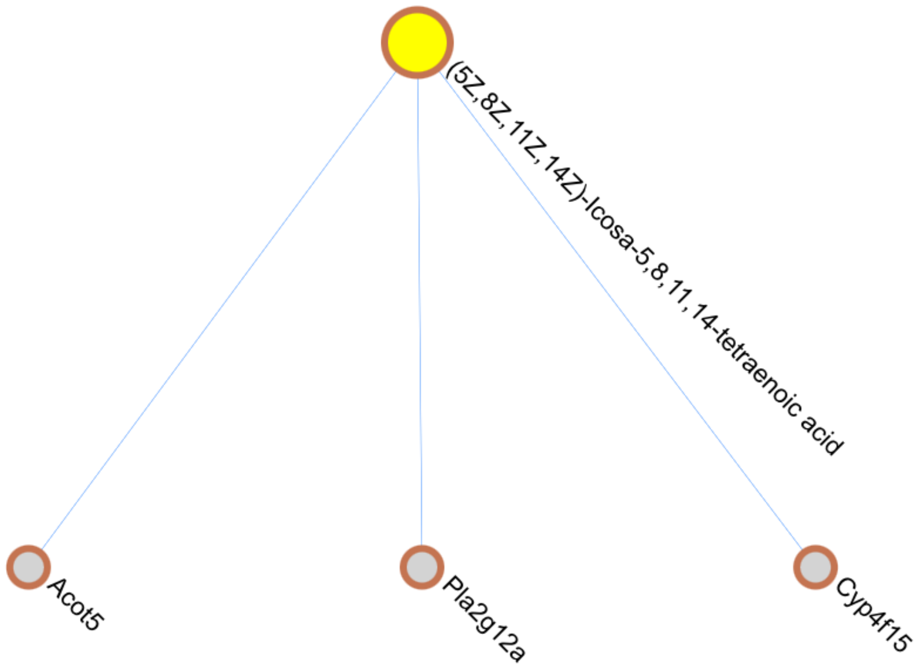
