## Supplementary material for "Multi-omics analysis in mouse primary cortical neurons reveals complex positive and negative biological interactions between constituent compounds in Centella asiatica": SF6.pdf

Module 2 – Fatty Acid Metabolism

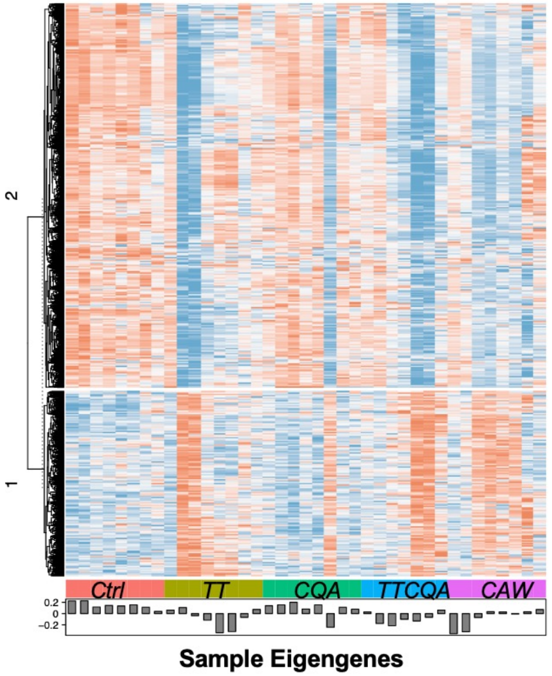


Module 3 – Cellular Response to Stress and Stimuli

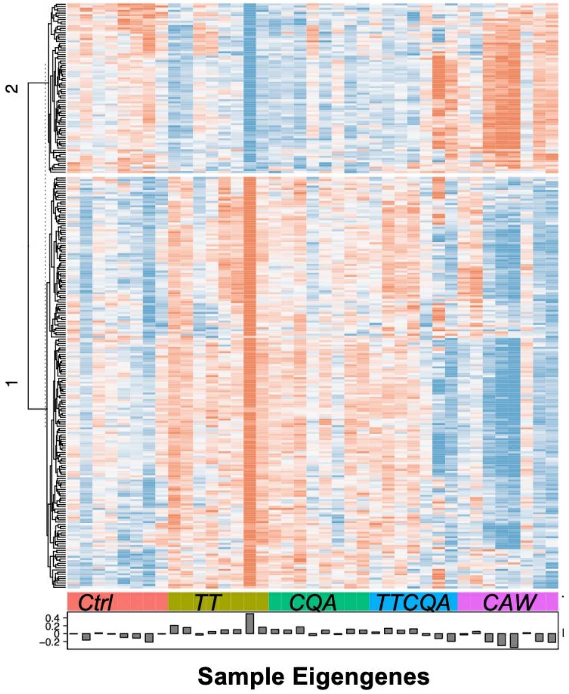


Module 4 – Immune Function

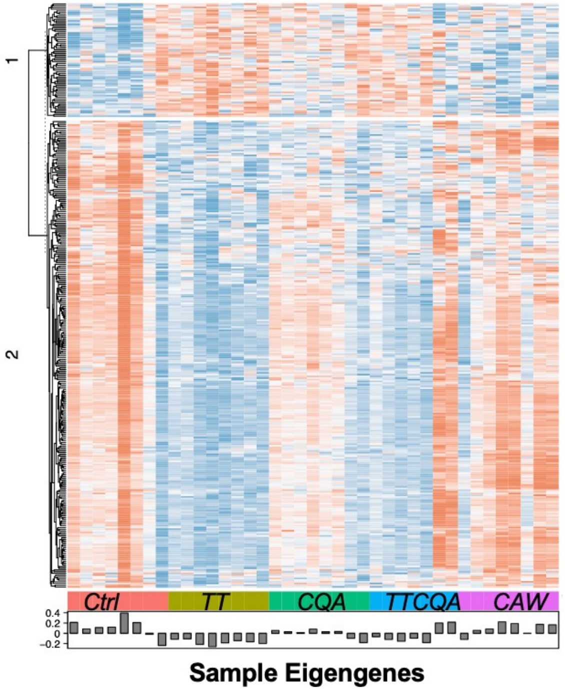


Module 5 – Electron Transport and Mitochondrial BioGenesis

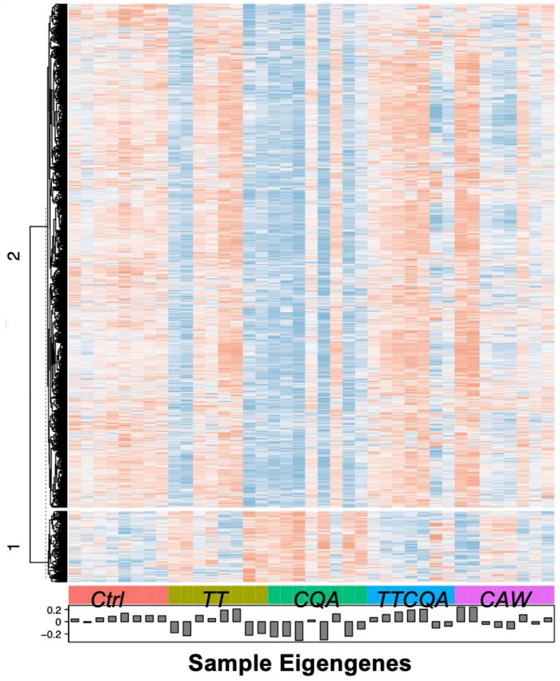
