## Supplementary material for "Multi-omics analysis in mouse primary cortical neurons reveals complex positive and negative biological interactions between constituent compounds in Centella asiatica": ST1.pdf

**Supplemental Table ST1.** Concentration (ng/mL) of phytochemical compounds in media cultured with neurons quantified using LC-MRM-MS.

The first four lines show expected starting concentrations (at time 0) of phytochemical marker compounds calculated from LC-MRM-MS analysis of concentrated stock solutions which were diluted 1/1000 into cell culture media. Phytochemical markers in 48h cultured media (CM) were measured in three biological replicates for each additive. All concentrations are reported as mean +/- 1 standard deviation of triplicate injections of each sample. Abbreviations: <LOD = less than limit of detection defined as 3 x standard deviation of the noise; <LOQ = less than limit of quantification defined as 10 x standard deviation of the noise; 5-CQA = 5-caffeoylquinic acid: 4-CQA = 4-caffeoylquinic acid; 3-CQA = 3-caffeoylquinic acid; 1,3-DiCQA =1,3-dicaffeoylquinic acid; 3,4-DiCQA = 3,4-dicaffeoylquinic acid; 3,5-DiCQA =3,5-dicaffeoylquinic acid; 1,5-DiCQA =1,5-dicaffeoylquinic acid; 4,5-DiCQA = 4,5-dicaffeoylquinic acid; MS = madecassoside; AS = asiaticoside; MA = madecassic acid; AA = asiatic acid.

| Sample ID | 5-CQA | 4-CQA | 3-CQA | 1,3-DiCQA | 3,4-DiCQA | 3,5-DiCQA | 1,5-DiCQA | 4,5-DiCQA | MS | AS | MA | AA |
|---|---|---|---|---|---|---|---|---|---|---|---|---|
| CAW Expected T0 | 173.0 ± 13.7 | 170.6 ± 10.1 | 418.8 ± 10.8 | 151.3 ± 12.7 | 134.4 ± 10.7 | 92.2 ± 8.3 | 202.2 ± 10.4 | 95.7 ± 1.6 | 1350.4 ± 53.4 | 611.6 ± 53.1 | 33.2 ± 25.7 | <LOQ |
| TT Expected T0 | <LOD | <LOD | <LOD | <LOD | <LOD | <LOD | <LOD | <LOD | 1280.0 ± 88.2 | 394.2 ± 18.3 | 15.1 ± 13.6 | <LOQ |
| CQA Expected T0 | 213.2 ± 17.6 | 197.2 ± 9.8 | 497.8 ± 25.0 | 130.3 ± 2.8 | 132.7 ± 3.9 | 109.6 ± 12.0 | 196.7 ± 26.7 | 87.5 ± 4.3 | <LOD | <LOD | <LOD | <LOD |
| TTCQA Expected T0 | 218.7 ± 29.9 | 203.1 ± 14.8 | 519.3 ± 32.4 | 140.4 ± 10.7 | 144.9 ± 10.6 | 113.6 ± 36.5 | 213.1 ± 27.4 | 84.5 ± 5.2 | 1433.3 ± 110.0 | 695.7 ± 38.3 | 5.0 ± 3.9 | <LOD |
| CAW7_CM_1 | 44.72 ± 0.30 | 30.10 ± 0.11 | 29.98 ± 0.23 | 14.38 ± 0.23 | 17.77 ± 0.33 | 7.21 ± 0.10 | 5.28 ± 0.22 | 13.09 ± 0.28 | 1865.46 ± 40.34 | 786.74 ± 8.76 | 66.89 ± 2.12 | 29.72 ± 0.38 |
| CAW7_CM_2 | 17.30 ± 0.18 | 9.0 ± 0.04 | 12.22 ± 0.18 | 2.16 ± 0.02 | 7.78 ± 0.10 | 3.69 ± 0.15 | 3.94 ± 0.02 | 4.88 ± 0.12 | 1933.35 ± 9.99 | 819.27 ± 1.27 | 65.63 ± 1.91 | 30.94 ± 3.96 |
| CAW7_CM_3 | 17.03 ± 0.04 | 8.87 ± 0.23 | 11.09 ± 0.21 | 3.24 ± 0.13 | 7.77 ± 0.20 | 3.80 ± 0.06 | 3.59 ± 0.03 | 5.28 ± 0.03 | 1930.66 ± 22.78 | 829.88 ± 7.67 | 65.67 ± 2.44 | 30.37 ± 4.67 |
| CQA_CM_1 | 221.20 ± 2.76 | 183.51 ± 2.08 | 169.95 ± 2.34 | 99.16 ± 1.09 | 76.86 ± 0.56 | 31.43 ± 0.69 | 18.15 ± 0.20 | 43.97 ± 0.34 | <LOQ | <LOQ | <LOQ | <LOD |
| CQA_CM_2 | 268.09 ± 5.99 | 225.89 ± 3.60 | 203.66 ± 3.08 | 156.17 ± 2.24 | 88.38 ± 0.95 | 39.93 ± 0.52 | 22.25 ± 0.07 | 47.57 ± 0.34 | <LOD | <LOD | <LOQ | <LOD |
| CQA_CM_3 | 234.85 ± 2.20 | 194.28 ± 3.08 | 184.06 ± 2.82 | 100.53 ± 1.07 | 86.92 ± 1.08 | 34.31 ± 0.55 | 20.68 ± 0.26 | 49.93 ± 0.79 | <LOQ | <LOQ | <LOQ | <LOD |
| TTCQA_CM_1 | 198.61 ± 1.07 | 167.30 ± 1.20 | 154.68 ± 1.61 | 98.15 ± 0.41 | 68.42 ± 0.86 | 28.79 ± 0.49 | 17.63 ± 0.15 | 38.69 ± 0.28 | 1627.16 ± 6.28 | 717.74 ± 10.99 | 21.27 ± 1.30 | 3.03 ± 0.13 |
| TTCQA_CM_2 | 304.3 ± 5.00 | 260.05 ± 5.74 | 231.44 ± 4.28 | 198.17 ± 4.31 | 101.95 ± 0.92 | 45.88 ± 0.93 | 22.88 ± 0.13 | 54.93 ± 1.00 | 1870.64 ± 38.16 | 830.79 ± 18.02 | 25.06 ± 2.27 | 4.49 ± 0.03 |
| TTCQA_CM_3 | 153.18 ± 3.47 | 125.71 ± 2.70 | 122.42 ± 1.44 | 63.25 ± 1.32 | 57.45 ± 1.06 | 22.76 ± 1.08 | 15.94 ± 0.42 | 33.20 ± 0.84 | 1605.32 ± 28.28 | 719.49 ± 11.76 | 27.74 ± 3.83 | 3.99 ± 0.38 |
| TT_CM_1 | <LOD | <LOD | <LOD | <LOD | <LOD | <LOD | 2.36 ± 0.07 | <LOD | 1620.88 ± 24.63 | 772.02 ± 10.20 | 21.08 ± 0.88 | 7.51 ± 0.39 |
| TT_CM_2 | <LOD | <LOD | <LOD | <LOD | <LOD | <LOD | 2.23 ± 0.05 | <LOD | 1675.60 ± 30.25 | 1027.18 ± 6.12 | 20.94 ± 2.31 | 9.50 ± 0.70 |
| TT_CM_3 | <LOD | <LOD | <LOD | <LOD | <LOD | <LOD | 2.28 ± 0.07 | <LOD | 1590.41 ± 45.15 | 734.17 ± 5.85 | 19.53 ± 0.45 | 7.62 ± 0.47 |
| MeOH_CM_1 | <LOD | <LOD | <LOD | <LOD | <LOD | <LOD | 2.18 ± 0.02 | <LOD | <LOD | <LOD | <LOQ | <LOD |
| MeOH_CM_2 | <LOD | <LOD | <LOD | <LOD | <LOD | <LOD | 2.29 ± 0.00 | <LOD | <LOD | <LOD | <LOQ | <LOD |
| MeOH_CM_3 | <LOD | <LOD | <LOD | <LOD | <LOD | <LOD | 2.30 ± 0.06 | <LOD | <LOD | <LOD | <LOQ | <LOD |
